# Poly(allylamine)/tripolyphosphate nanocomplex coacervate as a NLRP3-dependent systemic adjuvant for vaccine development

**DOI:** 10.1101/2024.07.01.601578

**Authors:** Gastón P. Rizzo, Rodrigo C. Sanches, Camila Chavero, Daiana S. Bianchi, Eugenia Apuzzo, Santiago E. Herrera, Maximiliano L. Agazzi, M. Lorena Cortez, Waldemar A. Marmisollé, Irene A. Keitelman, Analía S. Trevani, Sergio C. Oliveira, Omar Azzaroni, Paola L. Smaldini, Guillermo H. Docena

**Affiliations:** Instituto de Estudios Inmunológicos y Fisiopatológicos (IIFP), UNLP, CONICET, asociado a CIC PBA, Facultad de Ciencias Exactas, Departamento de Ciencias Biológicas, La Plata, Argentina (IIFP-UNLP-CONICET); Department of Biochemistry and Immunology, Institute of Biological Sciences, Federal University of Minas Gerais (ICB-UFMG).; Instituto de Investigaciones Fisicoquímicas Teóricas y Aplicadas (INIFTA), (UNLP, CONICET), 1900 La Plata, Buenos Aires, Argentina; Instituto de Química de los Materiales, Ambiente y Energía (INQUIMAE), UBA, CONICET, Facultad de Ciencias Exactas y Naturales, Departamento de Química Inorgánica Analítica y Química Física. Buenos Aires, Argentina; Instituto para el Desarrollo Agroindustrial y de la Salud (IDAS), (UNRC, CONICET), Ruta Nacional 36 KM 601, 5800 Río Cuarto, Argentina; Laboratorio de Inmunidad Innata, Instituto de Medicina Experimental (IMEX), CONICET, Academia Nacional de Medicina, Buenos Aires, Argentina; Department of Immunology, Institute of Biomedical Sciences, University of São Paulo (USP)

**Author notes:** Shared senior authorship (Paola L. Smaldini, Guillermo H. Docena). **Corresponding author:** Paola L. Smaldini.

**Keywords:** Coacervate, Nanoparticle, Adjuvant, Inflammasome, NLRP3, Systemic vaccines

## Abstract

Nanotechnology plays a crucial role in vaccine development. It allows the design of functional nanoparticles (NPs) that can act both as antigen carriers and as adjuvants to enhance the immune response. The present study aims to evaluate complex coacervate-like NPs composed of poly(allylamine hydrochloride) (PAH) and tripolyphosphate (TPP) as a safe vehicle and adjuvant for systemic vaccines. We investigated the activation of different antigen-presenting cells (APCs) with NPs and their adjuvanticity in Balbc/c and different KO mice that were intraperitoneally immunized with NP-OVA.

We found that NPs increased the expression of CD86 and MHCII and promoted the production and secretion of interleukin-1β (IL-1β) and IL-18 through the inflammasome NLRP3 when macrophages and dendritic cells were co-incubated with LPS and NPs. We evidenced an unconventional IL-1β release through the autophagosome pathway. The inhibition of autophagy with 3-methyladenine reduced the LPS/NPs-induced IL-1β secretion. Additionally, our findings showed that the systemic administration of mice with NP-OVA triggered a significant induction of serum OVA-specific IgG and IgG2a, an increased secretion of IFN-γ by spleen cells, and high frequencies of LT CD4^+^IFN-γ^+^ and LT CD8^+^IFN-γ^+^. Our findings show that NPs promoted the inflammasome activation of innate cells with Th1-dependent adjuvant properties, making them valuable for formulating novel preventive or therapeutic vaccines for infectious and non-infectious diseases.

**Graphical Abstract:** 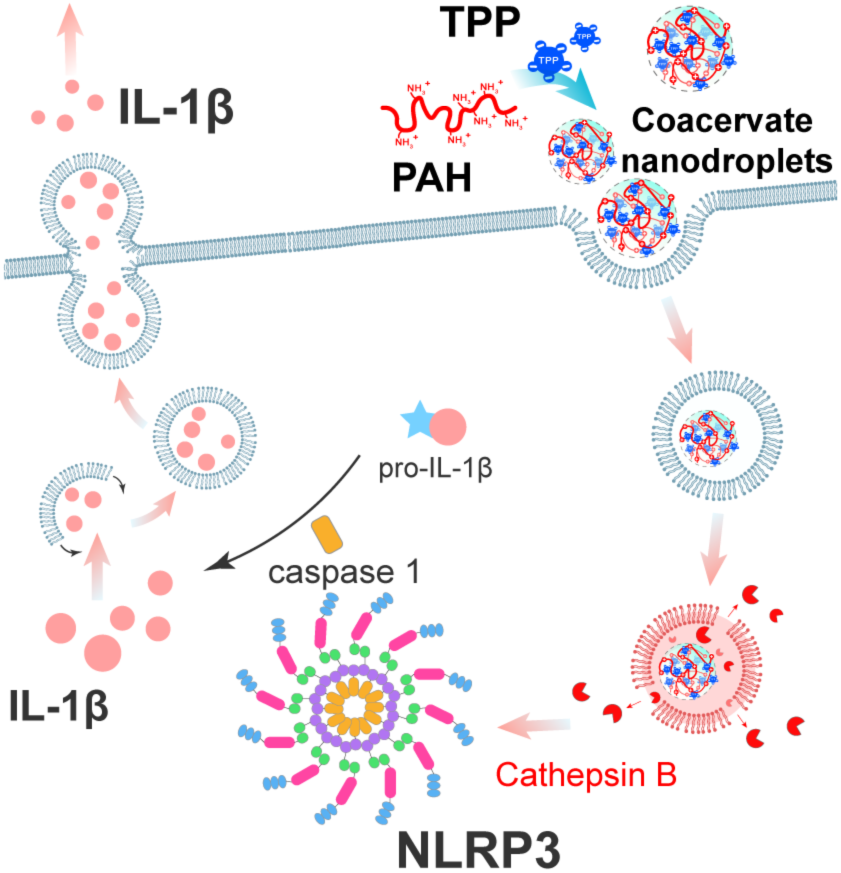

## INTRODUCTION

Vaccines have prevented millions of deaths in the last decades (mainly in the pediatric population), reduced medical care costs, and allowed us to coexist peacefully with pathogens.^1^ Protein antigen-based vaccines have emerged as attractive vaccine platforms as they are safer than traditional vaccines.^2^ However, they often generate a weak immune response when administered alone.^3^ To overcome this limitation, adjuvant agents are included in the formulations to enhance protection by induction of robust and perdurable immunity.^4^ These adjuvants trigger antigen-presenting cells (APCs) to release pro-inflammatory mediators and promote the clonal expansion of helper and cytotoxic T lymphocytes (CTLs), which target and destroy pathogen-containing bacteria and viruses, and tumor cells.^5,6^

Another limitation of protein subunit-based vaccines is that protein- or peptide-based antigens can be degraded by enzymes.^7^ Nanotechnology offers approaches for the design of nanostructures that allow encapsulation, protection and delivery of antigens.^3,8,9^ For instance, nanoparticle-based vaccines were shown to be safe and effective for immune intervention in COVID-19. ^10–12^ In addition, some nanosystems can enhance immune responses and function as self-adjuvants.^13–15^ Some nanoparticles have been designed to trigger innate immunity by activating the multiprotein complex known as the inflammasome in antigen-presenting cells.^16–18^ The process of internalization has been attributed to the similarities in size and morphological characteristics between nanoparticles and pathogens.^19,20^

Similar to the molecular patterns found in microorganisms (MAMPs and PAMPs) and signals from tissue damage (DAMPs) that activate the immune response, nanoparticle-associated molecular patterns (NAMPs) have been identified for non-pathogenic nanoparticles.^19,21^ NAMPs can interact with pattern recognition receptors on the surface or inside cells, initiating inflammatory responses that activate immune cells.^22,23^ Among these receptors, NLRP3, primarily expressed in myeloid cells like macrophages and dendritic cells, plays a key role in detecting NAMPs. When nanoparticles disrupt the lysosomal membrane, enzymes such as cathepsin B are released, triggering the activation of the NLRP3 inflammasome.^24^ Upon activation of the inflammasome, the adaptor protein ASC assembles into large aggregates, which recruit pro-caspase-1, leading to its activation. This process activates the pro-inflammatory cytokines IL-1β and IL-18 and the pore-forming protein gasdermin-D. Gasdermin-D pores facilitate the release of IL-1β and IL-18 while damaging the cell membrane, leading to an inflammatory form of cell death known as pyroptosis.^25,26^ However, some cytokines can be released through non-lytic pathways, including exocytosis or vesicle release, bypassing cell death.^27,28^ For instance, multivesicular bodies (MVBs), loaded with endosomes containing the cytokine, undergo lysis upon release into the extracellular environment, releasing the soluble cytokine.^29,30^ Additionally, alternative "non-canonical" inflammasome activation pathways involving caspase-11 have also been identified.^31,32^ Several delivery systems have been widely explored as adjuvant/delivery platforms for protein-based subunit vaccines, including liposomes, PLGA nanoparticles, micelles, and lipid−polymer hybrid nanosystems.^33–37^ Coacervation-like nanoplatforms based on ionic coacervation of polyamines (such as polyethyleneimine, chitosan, and polylysine) have demonstrated attractive properties as adjuvants.^38–40^ In general, coacervate-based nanoformulations have been designed to exhibit a positive surface charge as APCs more easily take them up with a negative surface charge.^7,41^ However, cationic systems usually present cytotoxicity due to their interaction with negatively charged biological membranes, which can impair cellular activity, such as membrane disruption.^42,43^ We recently reported anionic coacervate nanoparticles (size ∼200 nm and zeta potential ∼ −30 mV) produced through the direct and fully aqueous assembly of polyallylamine (PAH) with the tripolyphosphate anion (TPP), which exhibited long-term stability and ability to encapsulate proteins.^44^ Additionally, these nanostructures were non-toxic and demonstrated efficient uptake by APCs, such as macrophages.^45^

Given the promising properties of PAH/TPP nanoparticles, here we explored their potential as adjuvants to enhance immune system activation. Encapsulating an immunogen in this system may trigger the immune system by activating innate cells and endowing them with migratory capacity toward lymph nodes, where they can promote the activation of naïve B and T cells. This approach could make these nanoparticles suitable for vaccine formulation. Furthermore, activating these innate cells can lead to the production of cytokines and chemokines, along with the increased expression of key surface markers such as CD80, CD86, and MHCII. These changes transform the cells into potent APCs, enhancing their ability to activate T lymphocytes.^46–48^ In this work, we found that dendritic cells and macrophages undergo activation upon nanoparticle uptake and promotion of Th1-dependent adaptive immunity. The release of IL-1β and IL-18 through the NLRP3 activation (Figure 1A) triggered the secretion of IFN-γ with the induction of specific IgG antibodies (Figure 1B). Collectively, our findings indicated this PAH/TPP coacervate nanoformulation could serve a dual purpose: acting as a safe vehicle for antigen stabilization and delivery while also amplifying the immune response as nano adjuvants in the development of vaccine platforms.

**Figure 1.**
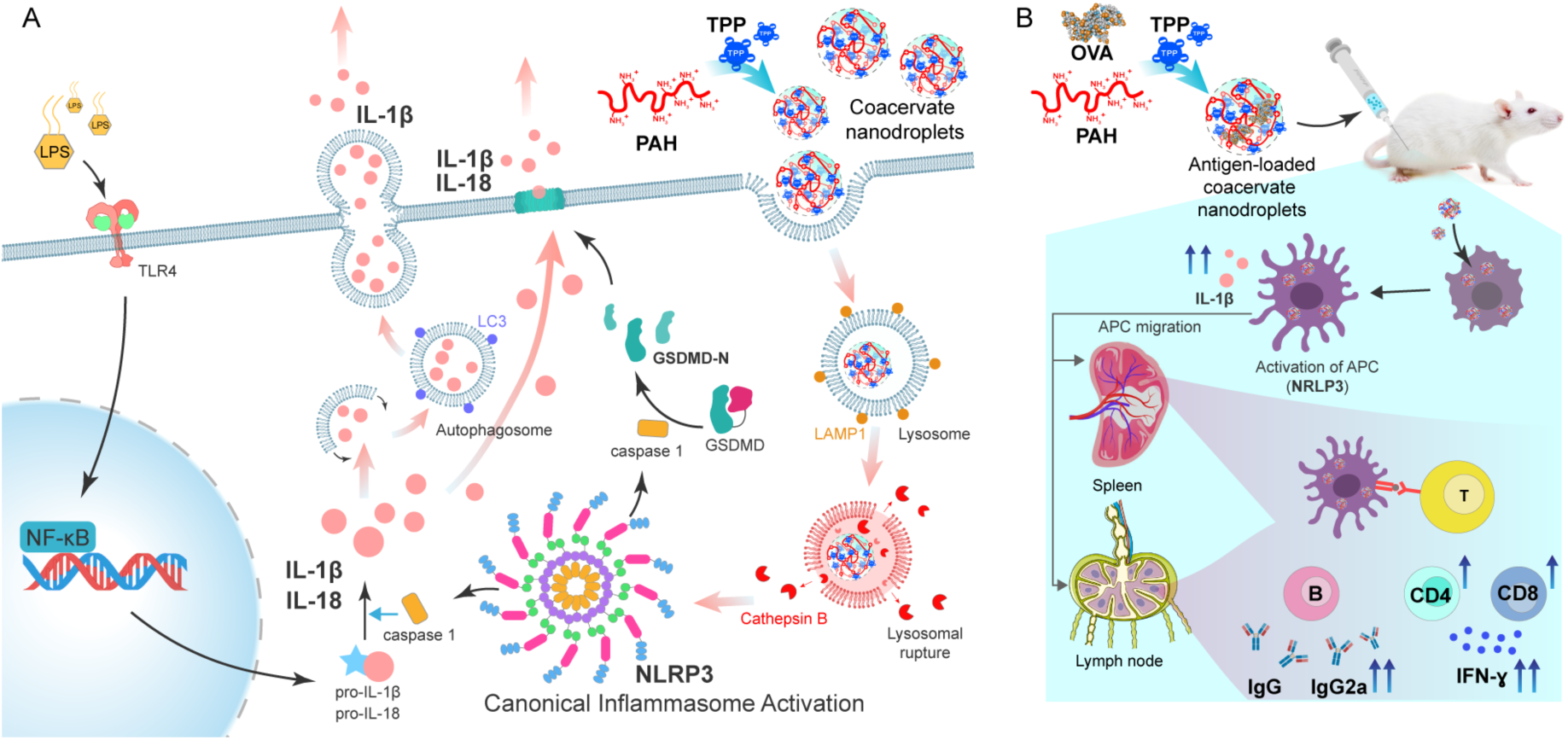
Mechanisms of NP-induced immune activation. **(A)** Schematic representation of NP uptake and NLRP3 inflammasome activation in phagocytic cells. Following NP internalization by phagocytosis, NPs are trafficked to lysosomes, where lysosomal membrane destabilization leads to the release of cathepsin B. Cathepsin B triggers the NLRP3 inflammasome pathway, leading to the activation of caspase-1, following the cleavage of Gasdermin D, and maturation of pro-IL-1β and pro-IL-18 for secretion. For the canonical inflammasome pathway, cells require an initial signal through Toll-like receptor to produce the pro-cytokines. Notably, the absence of pyroptosis, IL-1β can be released via the non-canonical autophagy pathway. **(B)** Systemic inoculation scheme with NP-OVA, leading to activation of phagocytic cells and subsequent migration to secondary lymphoid nodes, where they activate CD4+ and CD8+ T cells that produce IFN-γ and promote the activation of B cells for production of IgG and IgG2a antibodies.

## RESULTS

### Nanoparticle preparation and characterization

The interaction of polyamines with negatively charged multivalent small molecules was thoroughly studied over the past years. When mixing these components in an aqueous solution, a phase condensation takes place, leading to the formation of asymmetric polyelectrolyte complexes in which both intrinsic and extrinsic ion pairs are present between the long-chain polycation and short-chain polyanions.^49,50^ At charge-matching conditions (i.e. when the total amount of negative charges is equal to the total amount of positive charges), the mixing of polyamines with polyphosphates usually undergo macroscopic phases that go from insoluble solids (heavily crosslinked polyelectrolyte complexes) to polyelectrolyte-rich immiscible liquids (complex coacervates) depending on the pH and monovalent salt concentration. Recently, we reported the formation of an asymmetric coacervate composed of PAH (long-chain polyelectrolyte block) and TPP (small multivalent molecule).^44^ We showed that if the mixture is prepared under highly non-stoichiometric conditions and in the absence of added monovalent salts, it is possible to prevent the colloidal dispersion of coacervate droplets from collapsing into a macroscopic liquid phase, as typically occurs when symmetric coacervates are prepared under stoichiometric conditions. Figure 2A shows a simplified scheme of the preparation of a dispersion of nanometric-sized coacervate droplets formed by PAH chains ionically crosslinked by TPP anions (nanocomplexes). From here on, for simplicity, we will refer to these as nanoparticles (NPs).

**Figure 2.**
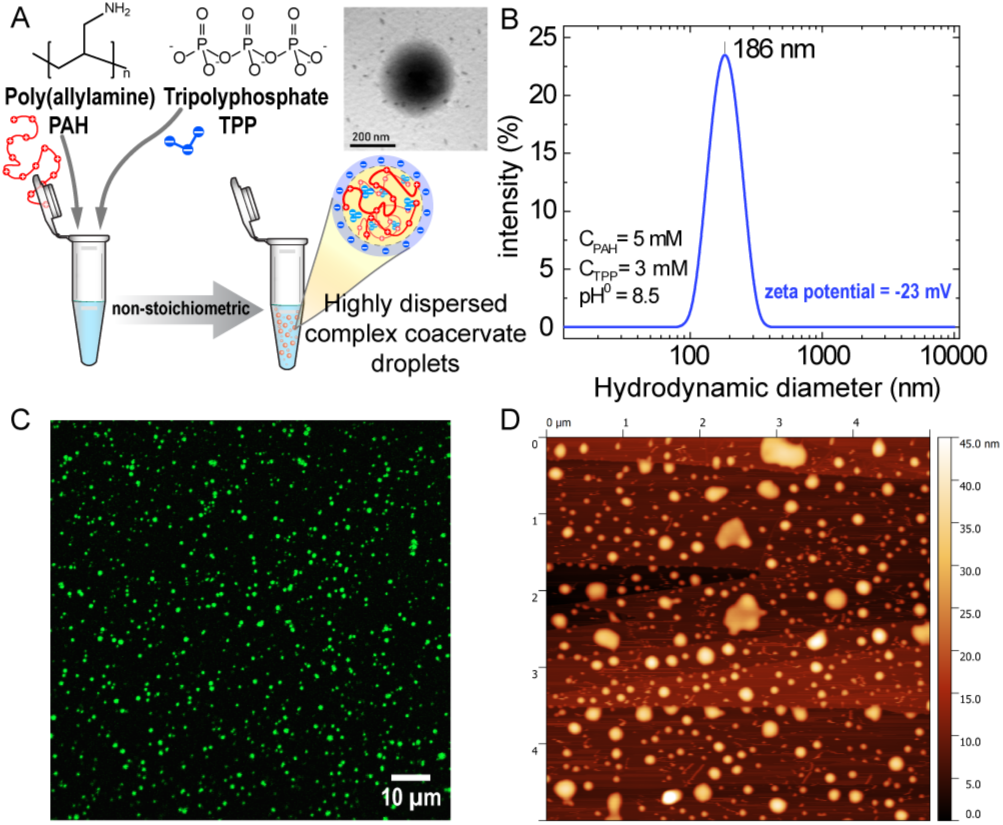
NP characterization. (A) PAH/TPP NP synthesis scheme. 5 mM PAH (monomer concentration) and 3 mM TPP are mixed in aqueous solution to obtain a highly stable colloidal dispersion of (asymmetric) coacervate droplets stabilized by electrostatic repulsions. Inset shows a TEM image of a single PAH/TPP NP. (B) DLS measurements showing a single size distribution (particle hydrodynamic diameter) centered at 186 nm. (C) Confocal microscopy image of a cover slip after being in contact over 2 min with a PAH*/TPP NP dispersion. (D) AFM (tapping mode) topography image of an HOPG flat substrate after being in contact over 2 min with a PAH/TPP NP dispersion.

Through dynamic light scattering (DLS) measurements, we detected a single distribution of particle sizes with a mean hydrodynamic diameter near 200 nm (Figure 2B and TEM) and a ζ-potential of −23 mV. Also, NPs were shown to remain stable in colloidal dispersion for more than 9 months after storing in a closed vessel,^44^ a key feature that can be explained by the negative surface charge of the NPs thanks to the excess of negative charges coming from the TPP anions. Also, we demonstrated that using a ratio of concentrations C_TPP_/C_PAH_=0.6, the excess of negative charges forces almost all of PAH chains (∼95%) to be ionically crosslinked, which means that no free PAH remains in solution phase.^44^ Using FITC-labeled PAH (PAH*), we observed that PAH*/TPP NPs exhibit a homogeneous size distribution when observed under confocal microscopy (Figure 2C). Finally, atomic force microscopy (AFM) measurements showed that, upon contact with a flat surface such as HOPG, NPs tend to deform by expanding in diameter and compressing in height, demonstrating their viscous nature (Figure 2D).

### Nanoparticle internalization triggers activation in phagocytic cells

To study the interaction of PAH/TPP NPs with cells, we exposed the murine J774 macrophages to NPs. We observed the internalization and cellular distribution of fluorescent NPs by confocal microscopy after 4h of incubation. Furthermore, we did not detect any internalization of the polymer PAH-FITC (without TPP) in macrophages (Figure S1). We observed that nanoparticles were well-dispersed into cytoplasmic vesicles (fluorescent granules) and the cytosol (diffuse fluorescence) (Figure 3A). Then, we analyzed the co-localization of the NP-FITC with LAMP1 to define the NP localization in vesicles. As it can be observed in Figure 3A, NPs were distributed between LAMP1^+^- and LAMP1^-^-vesicles, likely late and early endosomes (yellow and green vesicles, respectively). The incubation of cells along with the phagocytosis inhibitor cytochalasin D (Cyto D) confirmed that the internalization of NP was dependent on phagocytosis. Additionally, we detected lower fluorescence by flow cytometry in macrophages F4/80^+^ CD11b^+^ (Figure S1C) incubated with Cyto D than in cells exposed to NP-FITC (Figure 3B). Next, bone marrow-derived dendritic cells (BMDC) were exposed to NP or LPS and we analyzed the surface expression of Class II MHC and CD86 as cell activation markers (Figure 3C). We found that BMDC expressed significantly higher levels of Class II MHC and CD86 when exposed to LPS or NP than cells exposed to NPs or medium. The LPS used as a positive control for dendritic cell activation resulted in CD86 and Class II MHC expression similar to that in NP-exposed dendritic cells. These findings indicate that the cells recognized and were activated by PAH/TPP NPs.

**Figure 3.**
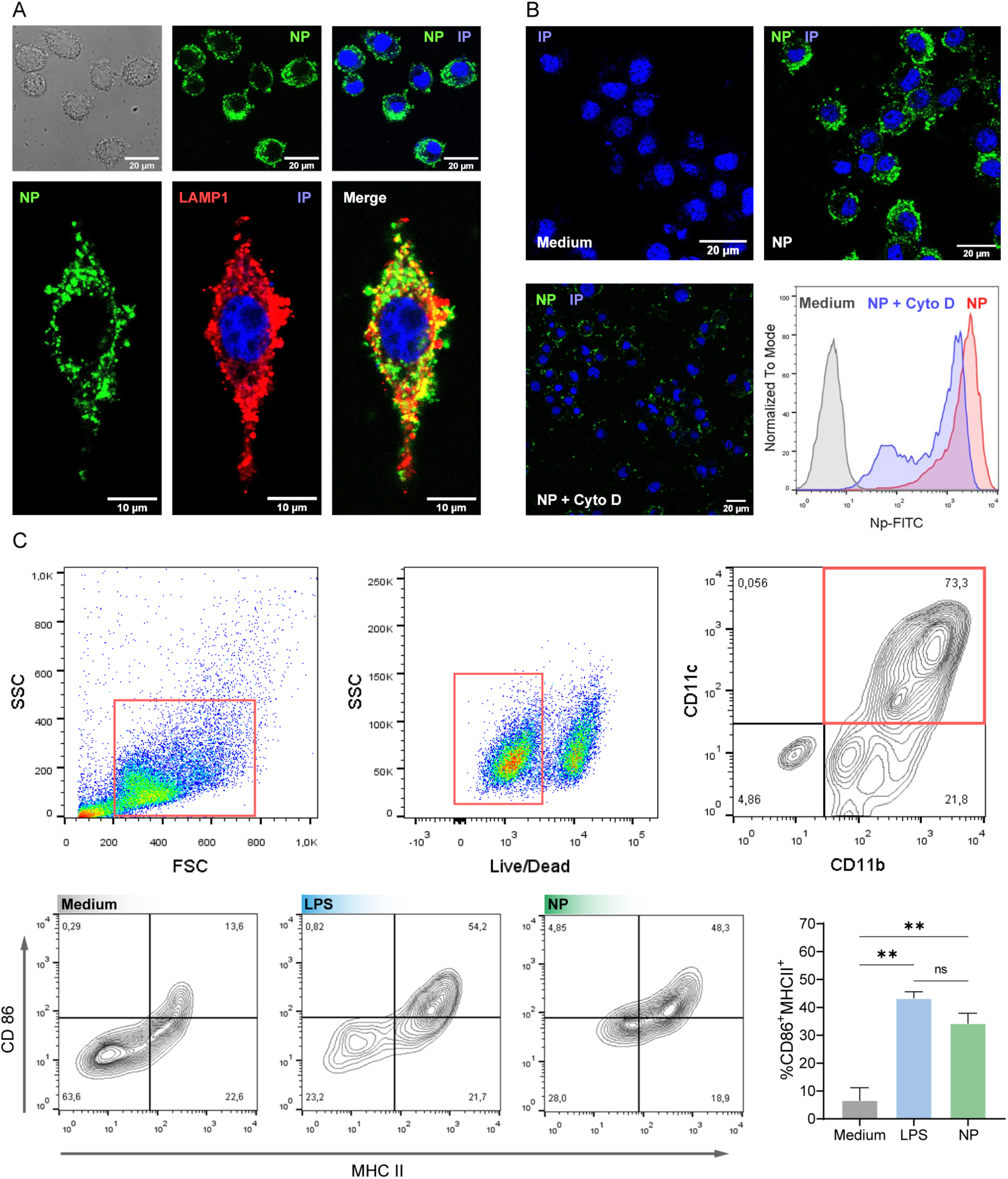
Cell uptake of NP and activation. **(A)** NP-FITC internalized by J774 and colocalized with the late endosome marker LAMP-1 by confocal microscopy after 4h stimulation. **(B)** Inhibition of phagocytosis of NP-FITC with cytochalasin D in BMDM by confocal microscopy and flow cytometry. **(C)** Surface expression of CD86 and MHCII in BMDC (CD11b^+^CD11c^+^) by flow cytometry. All experiments were performed in triplicate and data are expressed as mean±SEM. P-value was determined by ordinary one-way ANOVA *P<0,05; **P<0,01.

A similar analysis of internalization was conducted using a human intestinal epithelial cells line (HT-29). EPCAM was employed as a marker for epithelial cells, and various z-series were acquired by confocal microscopy. Neither intracellular fluorescence nor hIL-8 secretion was observed when epithelial cells were incubated with NPs. Pre-activated cells with TNF-α (20 ng/mL) rendered no NP uptake (Figure S2A) nor production of hIL-8 (Figure S2B). As a first approach to investigate the NP-driven secretion of mIL-1β in a phagocytic cell line, we performed an LPS (1 μg/mL) prime stimulus followed by a dose-response of NPs. We found that 0.5 mM NP (expressed as PAH monomer concentration) promoted the highest level of secreted mIL-1β (Figure S2C).

We next investigated the production of pro-inflammatory cytokines by phagocytic cells exposed to 0.5 mM NP or controls (Figure 4). We found a significantly higher level of IL-1β in murine J774 macrophages and BMDCs, and human THP-1 monocytes exposed to LPS+NP than cells exposed to LPS or NP (Figure 4A). We identified that the different cells exposed to NPs did not significantly increase IL-1β concentration compared with the medium; LPS+ATP was used as a positive control for IL-1β secretion.^51^ These findings collectively suggest that NP promoted IL-1β secretion and that LPS was necessary. Nevertheless, the quantification of secreted mIL-6 and hIL-8 showed no significant changes when cells were incubated with LPS or LPS+NP, indicating that the cytokine secretion was independent of the NP presence (Figure 4B). Similarly, the cytokine levels were basal when the cells were exposed to NP (Figure 4B). Furthermore, when we analyzed mIL-18, we observed a differential secretion in J774 macrophages and BMDC incubated with LPS+NP. Cells exposed to medium, LPS or NPs showed basal levels, whereas LPS+ATP or LPS+NP induced the highest concentration of mIL-18 in BMDC (Figure 4C). These findings indicate that the secretion of IL-1β and IL-18 in APCs required LPS and NP.

**Figure 4.**
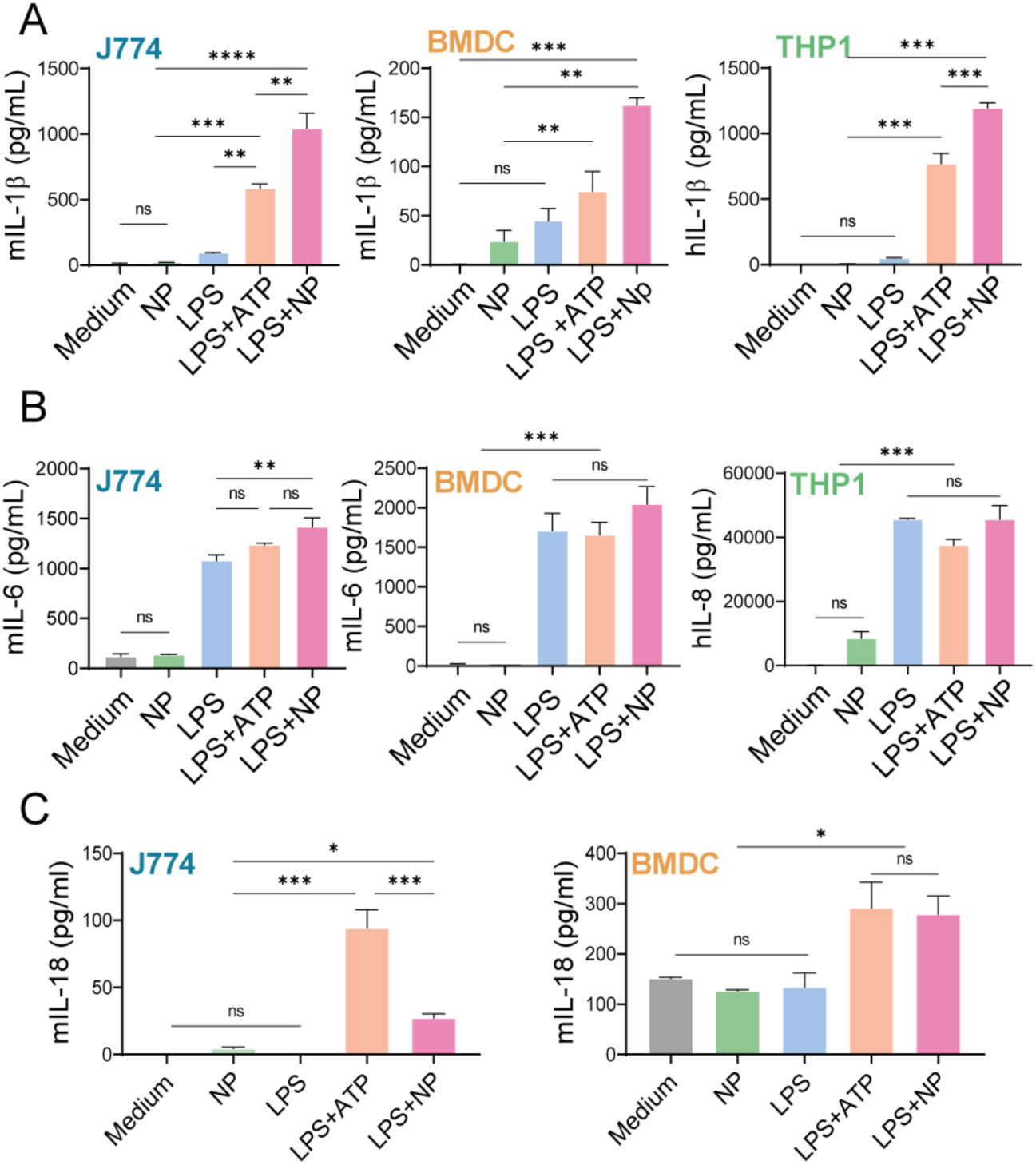
Secretion of pro-inflammatory cytokine by macrophages, dendritic cells and monocytes. J774, THP-1 and BMDC were prime with LPS (1μg/mL) for 3h; the medium was fully replaced and stimulated with ATP (50nM) for 3h as a positive control or with NP (0.5mM) overnight. Controls of LPS and NP were stimulated overnight. **(A)** IL-1β and **(B)** IL-6 and IL-8 and **(C)** IL-18 secreted cytokine by phagocyte cells were measured in supernatants by ELISA. All experiments were carried out in triplicate and data are expressed as mean±SEM. P-value was determined by ordinary One-way ANOVA *P<0,05; **P<0,01; ***P<0,001; ****p<0,0001.

### Nanoparticles activate the intracellular inflammasome pathway

Inflammasomes, such as NLRP3, require the adaptor molecule ASC to activate caspase-1.^52^ Following inflammasome activation, ASC assembles into a large protein complex, which can be visualized as a "speck".^53^ Representative pictures are shown with white arrowheads indicating GFP-specks (Figure 5A). The number of specks in LPS+NP-treated cells was significantly higher than in cells exposed to medium or LPS and equal to the positive control with LPS+DNA (p ≤0.05) (Figure 5B). This finding suggests that NP triggered the assembly of the ASC complex. A similar analysis was performed using flow cytometry to quantify ASC^+^ cells,^54^ and it was observed that cells exposed to LPS+DNA or LPS+NP had a significantly higher frequency of ASC^+^ cells compared to resting cells (Figure 5C and D). The ratio of GFP-ASC^+^ cells:total cells triggered by NPs was similar to the positive control condition of LPS+DNA. Additionally, the quantification of hIL-1β in the supernatants showed that LPS+DNA or LPS+NP induced higher levels of this cytokine than medium or LPS (p<0.05) (Figure 5E). To investigate the inflammasome activation, we used different pathway inhibitors. We found that cytochalasin D, the cathepsin B inhibitor CA-074Me and the caspase inhibitor Z-VAD-FMK significantly suppressed the mIL-1β secretion induced by LPS+NP (p<0.001) (Figure 5F). The analysis of the mIL-18 secretion showed similar results, with a higher secretion in cells exposed to LPS+NP than in cells incubated with LPS or medium, and the inhibitors suppressed the cytokine secretion (Figure 5F). Our results suggest that NP promoted inflammasome activation through cathepsin B and caspase-1. These findings led us to perform an experiment using inflammasome-null mice for NLRP3, caspase-1/11, caspase-11 and gasdermin D (GSDMD). BMDM obtained from NLRP3^-/-^, caspase1/11^-/-^, caspase11^-/-^ and GSDMD^-/-^ mice were incubated with 1μg/mL LPS and 0.5mM NP. To analyze the inflammasome activation, we quantified secreted mIL-1β and mTNF-α by ELISA in the supernatants. We observed no mIL-1β secretion in cells stimulated with LPS or NP in wild-type and KO mice. However, exposure to LPS+NP induced the production and release of this cytokine in wild-type, caspase11^-/-^ and GSDMD^-/-^ BMDM. In contrast, cells from NLRP3^-/-^ and caspase1/11^-/-^ mice showed decreased mIL-1β secretion compared to wild-type animals (p<0.001) (Figure 6A). When the BMDM WT and caspase11^-/-^ were stimulated with the positive control LPS+Nigericin, we detected the production of IL-1β. Furthermore, the secretion of IL-1β in BMDM from NLRP3^-/-^, caspase1^-/-^ and GSDMD^-/-^ was significantly abrogated (Figure 6A). The quantification of soluble mTNF-α showed that cells from wild-type and KO mice secreted similar high amounts upon exposure to LPS, LPS+NP or LPS+Nigericin. No TNF-α production was detected when cells were incubated with NP (Figure 6B). LDH measured in the supernatants showed that the stimuli did not significantly affect the cells. The consistent level of soluble LDH in cells exposed to the medium, NP or LPS+NP indicated that cell viability was unaffected (Figure S3A). Additionally, the lysate of LPS+NP-exposed cells from wild-type or KO mice showed a minimal difference in the activation of caspase-1 (20 kDa) between cells derived from wild-type and NLRP3^-/-^ mice. Inactive caspase-1 was detected in resting and LPS+NP-stimulated cells, while it was absent in samples from cells of caspase1/11^-/-^ animals (Figure S3B).

**Figure 5.**
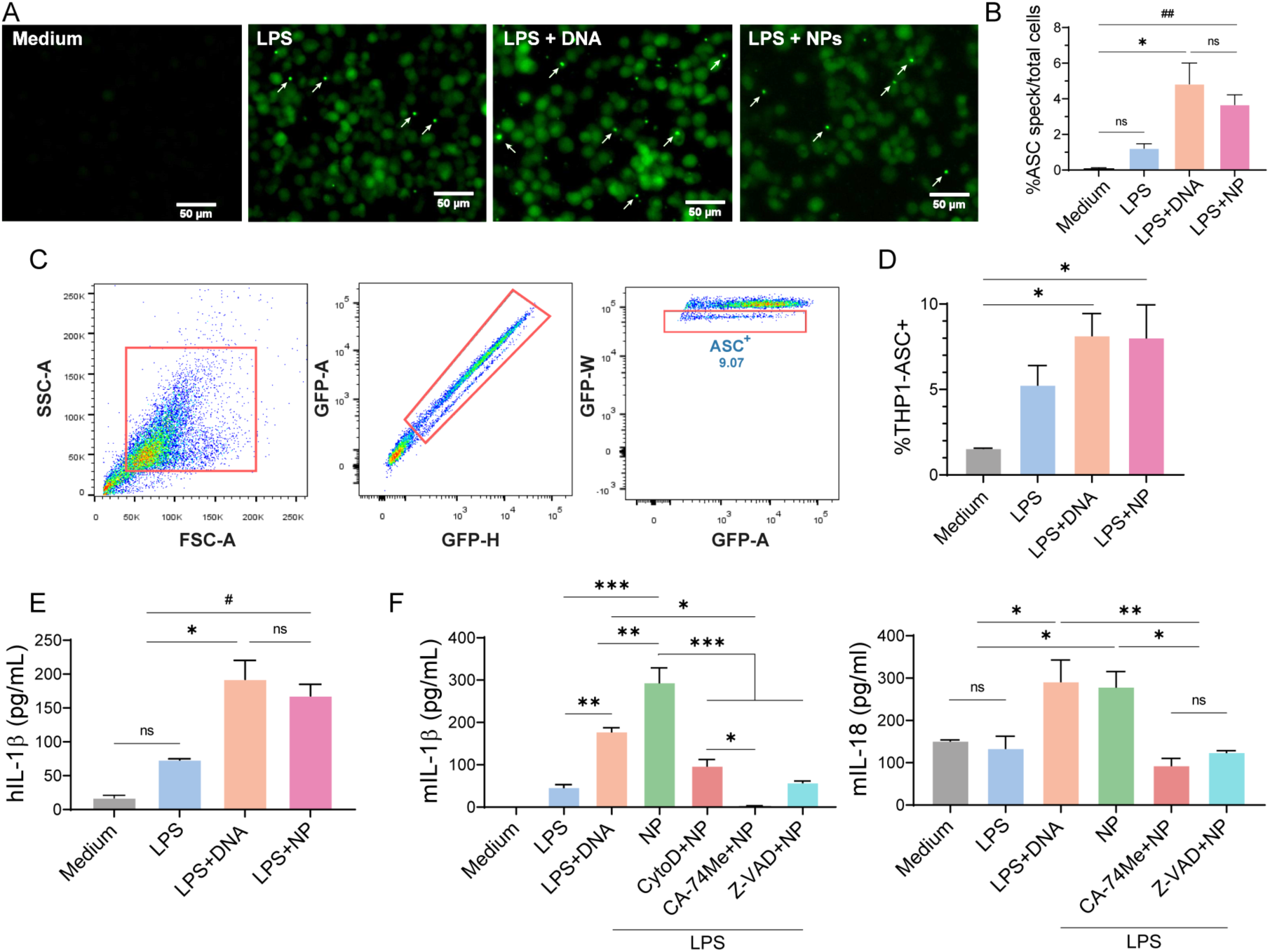
Inflammasome activation and cytokine secretion in THP1-ACS-GFP cells and BMDC. Cells were primed with LPS for 3 hours. The medium was fully replaced, and the cells were either stimulated with NP overnight or transfected with pcDNA as positive controls. **(A)** ASC**-**GFP specks within the cells are shown with a white arrow, performed by epifluorescence microscopy; **(B)** Frequency of speck ASC-GFP^+^cells. Three independent experiments were performed. Each bar represents the average number of total speck^+^ cells/total cells per field; 3 fields for each treatment were counted (2 replicates/treatment); **(C)** Flow cytometry gate strategy to define ASC-GFP^+^cells; **(D)** Frequency of speck ASC-GFP^+^cells by flow cytometry; **(E)** Quantification of secreted hIL-1β by ELISA. **(F)** BMDCs were pre-incubated with cytochalasin D (to inhibit phagocytosis), CA-74Me (to inhibit cathepsin B), and Z-VAD-FMK (Z-VAD; to inhibit caspases) for 30 minutes prior to overnight stimulation with LPS+NP. The levels of secreted mIL-1β and mIL-18 were quantified by ELISA. All experiments were carried out in triplicate and data are expressed as mean±SEM. P-value was determined by ordinary one-way ANOVA *p<0,05; **p<0,01; ***p<0,001 or t-Student #p<0,05; ##p<0,01.

**Figure 6.**
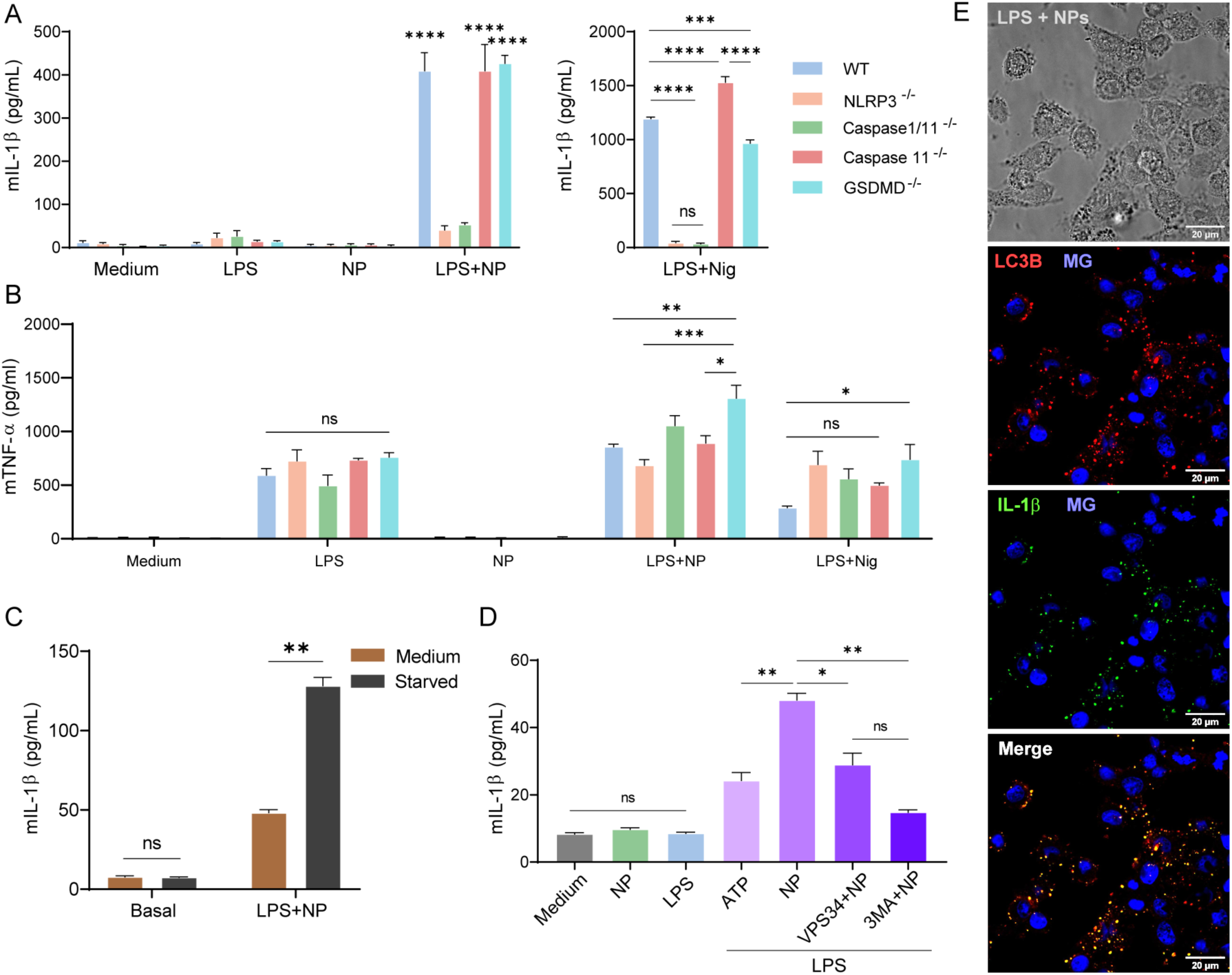
Inflammasome pathway analysis in BMDM from C57BL/6 (WT), NLRP3^−/−^, Casp1/11^−/−^, Casp11^−/−^, GSDMD^-/-^ mice. Cells were stimulated with LPS, NP, or LPS+NP, and **(A)** mIL-1β, **(B)** mTNFα were measured in supernatants by ELISA. The effect of autophagy stimulation by starvation on macrophage IL-1β secretion was investigated in J774 cell lines. After being primed for 3 hours with LPS, the cells were treated with NP. Six hours later, the cells were either left untreated in medium or had their supernatants replaced with Earle’s Balanced Salt Solution (EBSS) for an additional hour of culture and quantification **(C)** mIL-1β in the supernatants by ELISA. J774 cell line were stimulated for 6h with NP in the presence of inhibitors of the autophagy VPS34-IN1 and 3-methiladenine (3-MA) and quantification **(D)** mIL-1β. **(E)** Cytokine mIL-1β colocalized with LC3B by confocal microscopy after 6 hours of stimulation with LPS+NP (MG: Methyl green). Data are expressed as mean±SEM. P-value was determined by ordinary one-way ANOVA *P<0,05; **P<0,01; ***P<0,001; ****p<0,0001.

As mentioned before, we found that the release of mIL-1β was independent of GSDMD (Figure 6A). This prompted us to study the secretion of IL-1β. Among the multiple pathways described in myeloid cells, we investigated autophagy as responsible for the secretion of IL-1β. As starvation has been reported to induce autophagy,^55,56^ we performed experiments using the EBSS starvation medium following cell activation. We observed a significant increase in the production of mIL-1β when macrophages were stimulated with LPS+NP and then subjected to starvation with EBSS medium compared to the cells that were stimulated with LPS+NP (Figure 6C). To confirm this finding, we used 3-methyladenine (3-MA) (blocker of early stages of autophagy) and VPS34-IN1 (inhibitor of the class III PI3K) as autophagy inhibitors. All treatments significantly decreased the mIL-1β secretion in cells exposed to LPS+NP (Figure 6D). To confirm if autophagy is involved in the secretion of mIL-1β, we measured a pro-inflammatory cytokine (mIL-6) that is released through the canonical ER-Golgi and independent of the autophagy pathway. We observed unchanged high levels of mIL-6 in cells exposed to LPS+NP irrespective of the presence of EBSS (Figure S3C). However, when the different inhibitory conditions were assessed, we found that 3-MA led to a not significantly decreased cytokine level (Figure S3D). Nevertheless, it is essential to note that this effect was negligible compared to that exerted on mIL-1β secretion (Figure 6D). Additionally, we did not observe a significant increase in the release of LDH in the cells, either with or without the autophagy inhibitors (Figure S3E). Moreover, we observed similar results regarding IL-1β secretion when BMDC were treated with the autophagy inhibitors (Figure S3F). Finally, we found by confocal microscopy that mIL-1β co-localized with LC3B, a central protein in the autophagy pathway and autophagosome biogenesis, when macrophages were stimulated with LPS+NP (Figure 6E) compared to cells incubated with medium, NP or LPS (Figure S3G).

Overall, these findings confirmed that the NLRP3-dependent inflammasome pathway was involved in cell activation with IL-1β production, and that autophagy was involved in the secretion of IL-1β induced by LPS+NP. However, considering the reduced level of mIL-1β in cells from NLRP3^-/-^ mice compared with wild-type animals and the active caspase-1 observed in lysates of NLRP3^-/-^ cells (Figure S3B), we suggest that an NLRP3-independent inflammasome pathway may also be involved in mIL-1β production. Besides, the consistent level of soluble LDH in cells exposed to the medium, NP or LPS+NP indicated that cell viability was unaffected upon activation.

### The systemic administration of NP-OVA triggered a specific humoral and cellular immune response in mice

To characterize its adjuvanticity *in vivo*, we first measured the presence of LPS in PAH, TPP and NP with the human TLR4 HEK reporter cells, which secreted SEAP as readout. We found that the SEAP activity was comparable to that in non-stimulated cells and lower than 0.01 ng/mL of LPS (Figure S4). Next, we analyzed the ability of NPs to encapsulate proteins.

Considering previous results where we demonstrated that PAH/TPP nanoparticles can encapsulate a wide range of proteins such as lysozyme, cytochrome C oxidase, bovine serum albumin, and glucose oxidase,^44^ we tested the encapsulation of ovalbumin (OVA) as a model protein. The protein loading was carried out during the ionic crosslinking step in the synthesis of the NPs. Specifically, the protein was incorporated prior to the addition of TPP, forming PAH/OVA adducts that were then ionically crosslinked by the action of the TPP anion (Figure S5A). Although the protein encapsulation mechanism is not yet fully elucidated, we believe that proteins act as ionic crosslinkers to form three-component coacervates. In the case of OVA, whose isoelectric point is below physiological pH, it exposes negative surface charges, making it suitable as an ionic crosslinker for positively charged PAH. The analysis of OVA content in the supernatant after centrifuging the colloidal dispersion of PAH/OVA/TPP NPs (NP-OVA) showed that the encapsulation percentage is dependent on the OVA concentration used, with approximately 35% of protein encapsulated for 100 µg/mL OVA (Figure S5B).

To analyze the immune response, mice were intraperitoneally injected with NP-OVA in 3 weekly doses (Figure. 7A). As a control, we administered a formulation containing the approved adjuvant for human use, aluminum hydroxide (Alum),^57^ with OVA. No weight loss was observed during the whole experiment (Figure 7B). We did not observe specific antibodies to the nanoparticle or the polymer PAH in animals immunized with NP-OVA (Figure 7C). Animals that received NP-OVA produced higher levels of serum OVA-specific IgG than mice injected with OVA (p<0.01) and similar values compared with mice that received OVA+Alum (Figure. 7D). The subclass analysis revealed that NP-OVA induced higher titers of OVA-specific IgG2a than in mice injected with OVA+Alum (Figure 7E); in addition, the ratio of IgG subclasses in NP-OVA and OVA+Alum injected mice showed a higher production of OVA-specific IgG2a in NP-OVA immunized animals (Figure 7F). To analyze the T cell-mediated immune response, spleen cells from mice immunized with NP-OVA, OVA+Alum and OVA were stimulated with OVA and cytokines were measured in the supernatants. We found that mice immunized with NP-OVA induced increased levels of IFN-γ when cells were stimulated with OVA (p<0.001). Cells from mice that received OVA or OVA+Alum did not secrete IFN-γ (Figure 7G). On the other hand, the Th2 cytokines mIL-5 and mIL-13 showed decreased concentration in NP-OVA-treated mice compared to those administered with OVA+Alum (Figure 7G). To further evaluate the cell source of IFN-γ, spleen cells were analyzed by intracellular cytokine staining following *in vitro* OVA stimulation. We found that the frequency of CD4^+^IFN-γ^+^ and CD8^+^IFN-γ^+^ lymphocytes was significantly increased in mice immunized with NP-OVA (Figure 7H). Overall, these findings support that NP-OVA promoted the induction of a Th1-dependent immune response with specific IgG and IgG2a, and CD4^+^IFN-γ^+^ and CD8^+^IFN-γ^+^ T cells following the immunization with OVA adjuvanted with NP.

**Figure 7.**
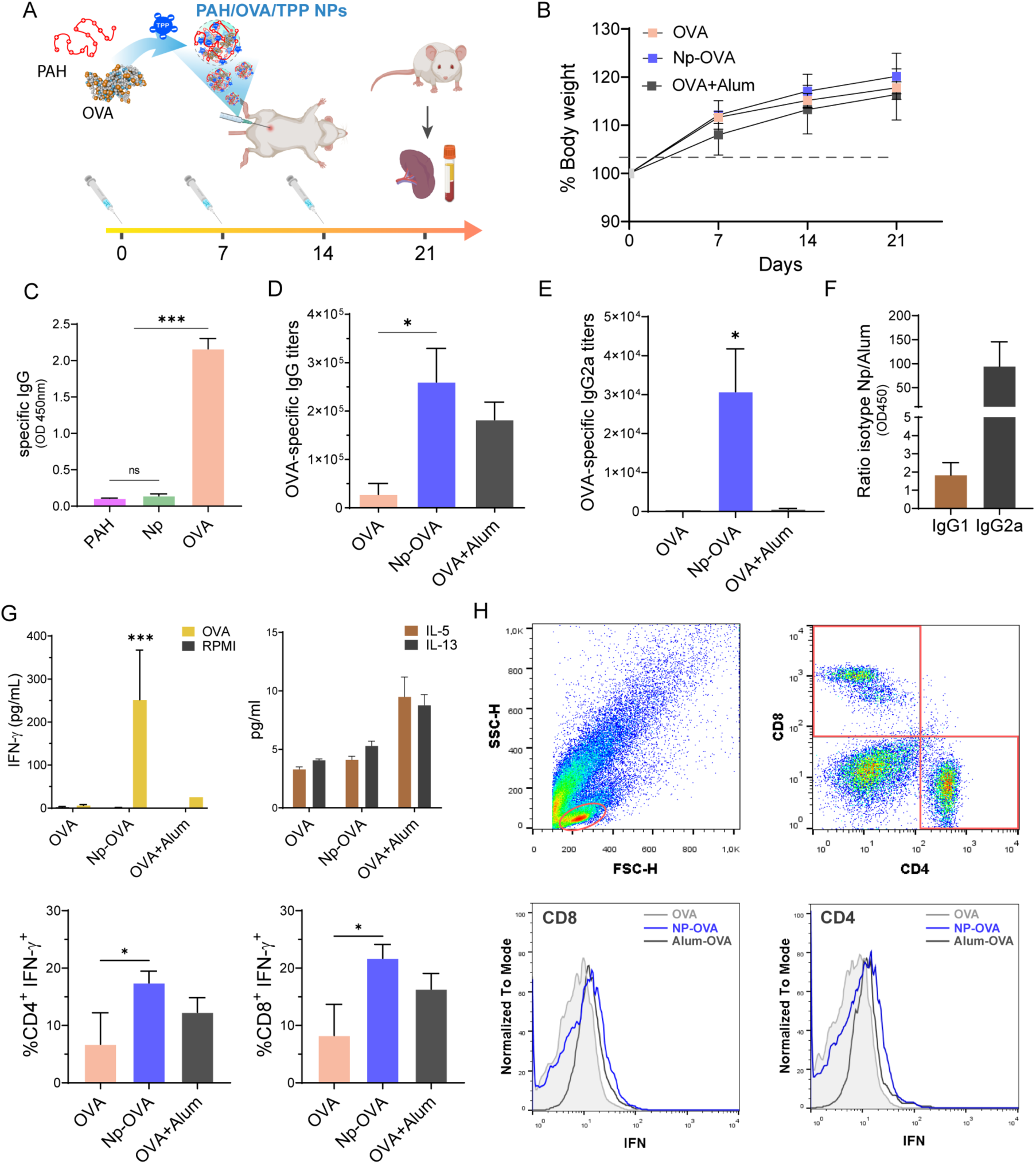
Intraperitoneally immunization of Balb/c mice with NP-OVA and specific humoral and cellular immune responses. **(A)** Immunization scheme; **(B)** Body weight monitoring; **(C)** quantification of serum specific IgG to PAH, NP and OVA and **(D)** OVA-Specific IgG and **(E)** IgG2a titers in serum of immunization animals and **(F)** serum optical density ratio of isotypes (IgG1 and IgG2a) between NP-OVA and OVA+Alum; **(G)** Quantification of mIFN-γ, mIL-5 and mIL-13 in the supernatants of spleen cells from immunized mice assessed by ELISA (top panel). Quantification for CD4^+^IFN-γ^+^ and CD8^+^IFN-γ^+^ T cells (lower panel). **(H)** Spleen cells from immunized mice were incubated with brefeldin A for the last 4h of culture and then were stained with anti-CD4 (PerCP-Cy5.5) or anti-CD8 (APC); flow cytometry gate strategy and Mean Fluorescent Intensity (MFI). The immunization experiment was carried out in duplicate, and all experiments were performed in triplicate; data are expressed as mean±SEM. P-value was determined by ordinary one-way ANOVA *p<0,05; **p<0,01; ***p<0,001 or t-Student test #p<0,05.

To investigate the role of the inflammasome pathway in promoting the specific immune response, C57BL/6 mice wild-type and NLRP3^-/-^, caspase1/11^-/-^ and IL-1R^-/-^ mice were intraperitoneally immunized with NP-OVA. As shown in Figure 8A, NLRP3^-/-^ and caspase1/11^-/-^ mice exhibited lower levels of serum OVA-specific IgG than wild-type mice (p<0.01), whereas IL1-R null mice showed similar levels than wild-type mice (Figure 8A). Finally, spleen cells from wild-type and KO mice were stimulated with OVA or medium, and IFN-γ levels were measured in the supernatants. We found that cells from wild-type mice immunized NP-OVA showed higher levels of IFN-γ than cells from WT mice that received OVA, as well as NLRP3^-/-^ and caspase1/11^-/-^ mice administered with NP-OVA. Additionally, cells from IL1-R^-/-^ mice injected with NP-OVA showed a similar concentration of IFN-γ to WT animals (Figure 8B). These findings indicate that the NLRP3 pathway is critical in promoting the OVA-specific Th1-dependent immune response.

**Figure 8.**
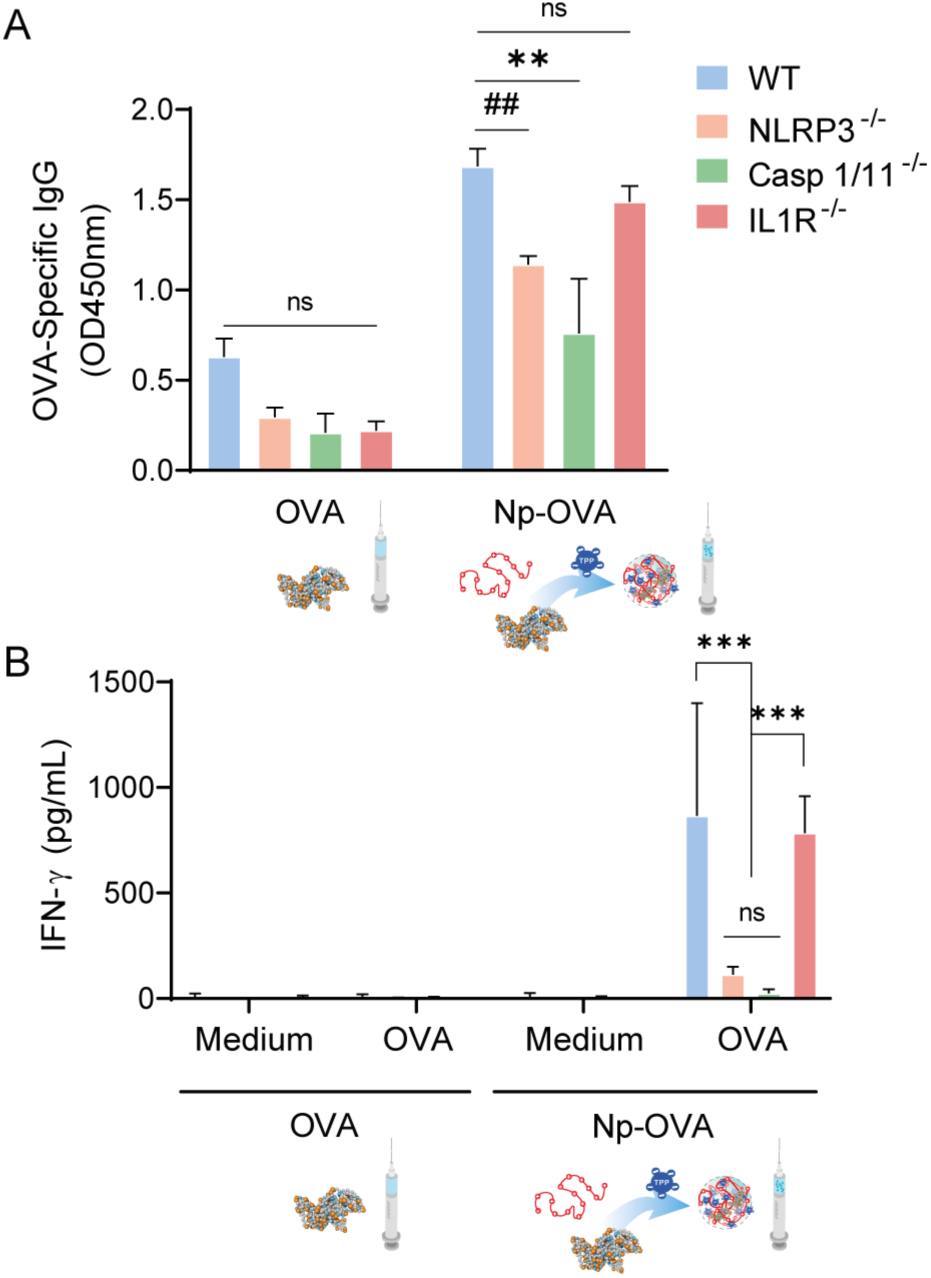
Intraperitoneal immunization with NP-OVA in WT and KO C57BL/6 mice and specific humoral and cellular response. C57BL/6 (WT), NLRP3^−/−^, Casp1/11^−/−^, IL-1R^−/−^ mice were immunized following the same scheme as in Figure 6A. **(A)** Measurement of serum OVA-specific IgG in serum (dilution 1/100) by ELISA and **(B)** secreted mIFN-γ by spleen cells from OVA or NP-OVA immunized mice by ELISA. The immunization experiment was carried out in duplicate and all experiments were performed in triplicate; data are expressed as mean±SEM. P-value was determined by ordinary one-way ANOVA *p<0,05; **p<0,01; ***p<0,001 or t-Student test ##p<0,01.

## DISCUSSION

Our findings showed that NPs are exclusively internalized by phagocytic antigen-presenting cells, specifically dendritic cells and macrophages. Subsequently, they localize intracellularly within the endosome compartment, causing their disruption and leading to the translocation of their components into the cytosol. Thus, the internalization of NP triggers the cellular activation and inflammasome assembly, releasing the pro-inflammatory cytokines IL-1β and IL-18 to the extracellular medium. Remarkably, we did not observe the NP internalization in non-phagocytic mucosa epithelial cells. The permeabilization of the lysosomal membrane is one of the most frequently described mechanisms for activating the inflammasome by particles and NP.^58,59^ We used several inhibitors to confirm the role of the different components of the pathway, and we corroborated the participation of cathepsin B as the early activator of the inflammasome.^60,61^ Then, using BMDM KO mice, we confirmed that cell activation followed by IL-1β secretion is NLRP3- and caspase-1-dependent and caspase-11- and gasdermin-D-independent with no induction of cellular death by pyroptosis.^62^ The literature shows that the non-canonical autophagy pathway is involved in the secretion of inflammatory molecules such as IL-1β and HMGB1.^63,64^ In macrophages and monocytes, LC3B is essential for the efficient secretion of IL-1β in response to lysosomal damage.^56,65^ In this sense, we demonstrated that the inflammasome activation proceeded with autophagy induction for secreting IL-1β. All together, these findings demonstrated that PAH-based NP activated phagocytic cells with increased expression of MHC II and CD86, along with the secretion of IL-1β and IL-18, without inducing cell death, which make them a candidate adjuvant to trigger an antigen-specific immune response.

It is known that aluminum salts, such as the extensively used in human vaccines aluminum hydroxide (alum), can enhance the effectiveness of the immune response by activating antigen-presenting cells through the depot effect and the aggregation of antigens to improve the uptake and activation of APCs. Furthermore, it has been described that alum triggered the NLRP3-dependent inflammasome pathway to further promote adaptive immunity and memory.^66^ Thus, we evaluated the *in vivo* adjuvant capacity of PAH/TPP NP in mice that were systemically administered with OVA encapsulated in NP. We observed the induction of innate immunity via the inflammasome-dependent pathway to further promote a Th1-driven adaptive humoral and cellular immunity. Our findings showed the induction of serum IgG and IgG2a-specific antibodies and the expansion of CD4^+^- and CD8^+^-IFN-γ-producing T cells. Furthermore, we confirmed *in vivo* that the Th1-dependent isotype synthesis and IFN-γ production were NLRP3- and caspase-1-dependent. It is reported that the NLRP3 inflammasome pathway initiates a caspase-1–dependent IL-1β and IL-18 secretion promoting interferon-γ production and Th1 differentiation.^67,68^ Similar to other adjuvants, such as Alum and QS-21, PAH-based nanoparticles, along with LPS as a first inflammasome activation signal, induced *in vitro* the NLRP3-ASC-caspase-dependent IL-1β production. *In vivo*, however, the first signal may be triggered by a DAMP, such as uric acid, generated at the immunization site.^69,70^ Additionally, IL-18, constitutively expressed by several cell types, can stimulate IL-1β transcription and may initiate a pro-inflammatory cascade during vaccination.^71^ The *in vivo* induction of OVA-specific CD8^+^T cells indicates that NP-OVA triggered the cross-presentation of OVA, which may proceed upon the endosome is disrupted and the content is translocated into the cytosol. Further studies are needed to confirm if macrophages and dendritic cells contribute to this mechanism, which is critical when considering using PAH-based NP as an adjuvant in an anti-tumoral or viral vaccine.

Muñoz-Wolf et al. reported that polymeric nanoparticles made of poly-lactic-co-glycolic acid with a diameter of 50-60nm elicited a robust cellular immune response through the caspase-11- and the pyroptosis effector gasdermin D-dependent non-canonical inflammasome pathway. They described a division of labor for IL-1β and IL-18 to induce the appropriate immunogenic context for Th1 and CD8^+^ T cell induction, respectively.^72^ In our study, we characterized a polymeric nanoparticle of PAH that activated murine and human macrophages and dendritic cells through the canonical caspase-1- and NLRP3-dependent inflammasome pathway with the secretion of IL-1β and IL-18, without cell death, and independent of caspase-11 and gasdermin D-dependent pyroptosis^73,74^. We found that PAH-based NP induced autophagy for the secretion of the pro-inflammatory cytokine IL-1β.

## CONCLUSION

In summary, we characterized a novel polyallylamine-based self-assembled nanoparticle as a systemic and safe adjuvant, demonstrating its ability to target the encapsulated OVA immunogen to phagocytic cells efficiently. This targeting enabled antigen-presenting cells to acquire the capacity to activate T cells, thereby inducing a robust Th1-dependent humoral and cellular immune response. The ability of this nanosystem to enhance the immune system’s recognition and response to the antigen highlights its potential for biomedicine. These findings advance our understanding of nanoparticle-based adjuvants and pave the way for designing and developing innovative nanoparticle-based systemic vaccines, which could be used to treat a wide range of infectious and non-infectious diseases, potentially transforming vaccine strategies in immunotherapy.

## MATERIALS AND METHODS

### Preparation and characterization of PAH/TPP nanoparticles

PAH/TPP nanoparticles were prepared by following the procedure described in Apuzzo et al^45^. Briefly, 0.125 *V_f_* ml of PAH (17.5 kDa, Sigma-Aldrich, St. Louis, MO, USA) 40mM (monomer concentration) at pH 8.5, 0.75 *V_f_* ml of deionized water, and 0.125 *V_f_* ml of TPP (Sigma-Aldrich, St. Louis, MO, USA) 24mM at pH 8.5 were mixed under constant stirring at room temperature (RT). The addition of TPP was made very quickly to prevent the system from collapsing into a macroscopic phase. After this procedure, a volume *V_f_* of dispersed PAH/TPP nanoparticles is obtained with a final concentration of PAH and TPP of 5mM and 3mM, respectively. In this work, the NP concentration was expressed in terms of PAH monomer concentration (i.e., 0.5mM NP means a 1:10 dilution of the original dispersion). The protein-loaded nanoparticle dispersions were prepared by following the same procedure but adding an aqueous solution of chicken egg albumin (OVA) (EndoFit^(TM)^ InvivoGen, San Diego, CA, USA) at the water addition stage. The volume of deionized water was re-calculated as *V_water_* = 0.75 *V_f_* − *V_OVA_* and the OVA volume was fixed to obtain the desired OVA concentration in the final solution. The percentage of encapsulated OVA was calculated by assessing the protein content of the supernatant using bicinchoninic acid (BCA) commercial kit (Thermo Fisher Scientific, Waltham, MA, USA).

Atomic force microscopy (AFM) images were taken with an Agilent 5500 SPM in tapping mode at room temperature using PPP-NCL (NANOSENSORS^TM^) silicon tips. For the sample preparation, a droplet of the suspended PAH/TPP NPs was placed on top of a freshly cleaved highly oriented pyrolytic graphite (HOPG) substrate, rested for 2 min, rinsed with Milli-Q water, and dried with N_2_. Transmission electron microscopy (TEM) images were acquired with a JEOL microscope (120 kV) equipped with a Gatan US1000 CCD camera. For the TEM sample preparation, 5 µl of a PAH/TPP NPs dispersion was placed in a carbon-coated copper grid and stained with phosphotungstic acid to create image contrast. Dynamic light scattering (DLS) and ζ-potential measurements were performed using a ZetaSizer Nano (ZEN3600) at 20 °C using disposable cuvettes. Hydrodynamic diameter was measured using a 173° backscatter angle with 20 runs per sample, averaging 5 measurements. ζ-potential was calculated from electrophoretic mobility with 100 runs per sample. Confocal (epifluorescence) microscopy of PAH/TPP NPs was carried out using a Carl Zeiss Axio Observer 7 with a Plan-Apochromat 63x/1.40 Oil DIC M27 objective.

### Cell lines

HEK-hTLR4-reporter (InvivoGen®, San Diego, CA, USA), HT-29 and J774 cell lines were maintained in DMEM medium (Gibco® Thermo Fisher Scientific, Waltham, MA, USA) containing 10% FBS (Internegocios, Mercedes, Bs As, Argentina), 100U/mL penicillin and 100µg/mL streptomycin (Gibco® Thermo Fisher Scientific, Waltham, MA, USA). The THP1-ASC-GFP cell line was maintained in RPMI medium (Gibco® Thermo Fisher Scientific, Waltham, MA, USA) containing 10% FBS, 100U/mL penicillin and 100µg/mL streptomycin.

Bone marrow-derived macrophages (BMDMs) were cultured in DMEM medium, and bone marrow-derived dendritic cells (BMDC) were cultured in RPMI medium containing 10% FBS, 100U/mL penicillin and 100µg/mL streptomycin. Both cells were derived from wild-type (WT) and knockout (KO) mice with 20% L929-conditioned medium (LCCM) in DMEM and 20ng/mL recombinant granulocyte-macrophage colony-stimulating factor (GM-CSF) (R&D Systems, Minneapolis, MN, USA) in RPMI for BMDM and BMDC, respectively. Briefly, bone marrow cells were obtained from the femur and tibia of WT or KO mice^75^. Cells were seeded on dish plate 5×10^5^ cell/mL (day 0) and maintained in a conditioned medium and GM-CSF containing 10% FBS, 100U/mL penicillin and 100µg/mL streptomycin at 37°C, in a 5% CO_2_ atmosphere for 7 days. On day 4, the medium was fully replaced, and cells were stimulated with 0.5mM PAH/TPP NPs. As a positive control for inflammasome activation, cells were treated with 5mM Nigericin for 50min, after being primed with 1μg/mL LPS from *E. coli* (LPS E. coli) (Sigma-Aldrich, St. Louis, MO, USA) for 3h at 37°C. Supernatants were collected and cells were lysed with 25µL/well M-PERR (Thermo Fisher Scientific, Waltham, MA, USA) supplemented with 1:100 protease inhibitor cocktail (Sigma-Aldrich, St. Louis, MO, USA). Both supernatants and cell lysates were stored at −80°C until used.

### Cell stimulation with NP and characterization

#### Flow cytometry for phenotype and cell activation characterization

BMDC were seeded in 24-well microplates at 1×10^6^ cells/well and stimulated with ultrapure 1µg/mL LPS from *E. coli* for 3h at 37°C, followed with 0.5mM PAH/TPP NPs overnight (ON). BMDC were washed with PBS and stained for 1h at 4°C with anti-CD11c (PE), anti-CD11b (PerCP-Cy5.5), anti-MHCII (APC) and anti-CD86 (FITC) (Thermo Fisher Scientific, Waltham, MA, USA). As a positive control for inflammasome induction, cells were primed with ultrapure 1µg/mL LPS from *E. coli* for 3h, followed by 5mM ATP (Sigma-Aldrich, St. Louis, MO, USA) for 3h at 37°C. For inhibition analysis, cells were pre-treated for 30min with 100µg/mL Z-VAD-FMK (Sigma-Aldrich, St. Louis, MO, USA), 1µg/mL cytochalasin D (Santa Cruz Biotechnology Inc, Santa Cruz, CA, USA) or 50µM CA-74ME (Santa Cruz Biotechnology Inc, Santa Cruz, CA, USA) and kept in culture during the experiment. Fluorescence acquisition was performed with a FACS Aria Fusion cytometer, and data was analyzed using FlowJo software.

#### Reagents for inhibitions

For inhibition analysis, cells were pre-treated for 30min with 100µg/mL Z-VAD-FMK, 1µg/mL cytochalasin D or 50µM CA-74ME, 5mM 3-methyl adenine (Cayman, Ann Arbor, MI, USA) and 1 µM VPS34-IN1 (Cayman, Ann Arbor, MI, USA) and 100nM Bafilomycin A1 (Cayman, Ann Arbor, MI, USA) and kept in culture during the experiment. When indicated, at 4 hours post-NP stimulation, supernatants were removed and replaced with either complete medium or Earle’s Balanced Salt Solution (EBSS) for medium starvation.

#### Nanoparticle internalization analysis

BMDC and BMDM were grown on glass coverslips and stimulated ON with 0.5mM FITC-PAH/TPP NP. Poly (fluorescein isothiocyanate allylamine hydrochloride), Mw∼56000 (Sigma-Aldrich, St. Louis, MO, USA) was used instead of PAH in the PAH/TPP formulation. Cells were fixed for 30min in 4% paraformaldehyde (PFA) at RT and nuclei were stained with 1µg/mL propidium iodide (IP) (Sigma-Aldrich, St. Louis, MO, USA). Cells were washed with PBS, blocked with 2% BSA in phosphate buffer pH=7.4 for 30min at RT and incubated with mouse anti-human LAMP1 (BD Biosciences, Franklin Lakes, NJ, USA) for 1h at RT, followed with anti-mouse Alexa 647 (Abcam, Cambridge, UK) for 1h at RT. Cells were washed, coverslips were mounted onto a slide containing 10µL of fluorescence mounting media (Dako, Abcam, Cambridge, UK) and allowed to dry at 4°C ON. Fluorescence was analyzed with a Leica SP5 Confocal Microscope.

#### Internalization inhibition assay

BMDC and J774 were stimulated with 0.5mM FITC-PAH/TPP NPs and phagocytosis was inhibited with 1µg/mL cytochalasin D. Cells were characterized by flow cytometry with anti-CD11b (Percp-Cy5.5) and anti-F4/80 (APC) (Thermo Fisher Scientific, Waltham, MA, USA) and confocal microscopy for intracellular fluorescence. Nuclei were stained with 1µg/mL IP.

#### Intracellular immunostainings for the localization of IL-1β in autophagosomes

*The* J774 cell line was grown on glass coverslips and prime for 3h at 37°C with 1µg/mL LPS from E. coli, followed with 0.5mM PAH/TPP NP for 6h. After fixation with PFA 4% for 30 min, cells were blocked with PBS-glycine (0.1 M) for 15 min, permeabilized with acetone (−20°C) for 7min, and blocked with PBS-BSA 5% for 1h at RT. Then, macrophages were incubated with the rabbit polyclonal anti-LC3B (Proteintech, Rosemont, IL, USA) and mouse polyclonal anti-IL-1β (Cell Signaling Technology, Danvers, MA, USA) in blocking buffer ON at 4°C and then incubated with the secondary anti-rabbit Alexa 594 (Abcam, Cambridge, UK) and anti-mouse Alexa 488 (Invitrogen, Thermo Fisher Scientific, Waltham, MA, USA) antibodies for 1h at RT. Nuclei were stained with methyl green (Sigma-Aldrich, St. Louis, MO, USA) and mounted onto a slide containing fluorescence mounting media (Dako, Abcam, cambrige, UK). Image acquisition was performed with an SP5 Leica Confocal Microscope.

#### Epithelial cell stimulation

HT-29 cells were maintained in DMEM containing 10% FBS, 100U/mL penicillin and 100µg/mL streptomycin at 37°C with 5% CO_2_, and 3×10^4^ cells/well were seeded in 96-well microplates. Cells were incubated ON to 70% confluence and the medium was fully replaced. Then, cells were stimulated ON with 0.5mM FITC-PAH/TPP NP, or with 50ng/mL TNF-α (R&D Systems, Minneapolis, USA) for 30min at 37°C followed with FITC-PAH/TPP NP. A mouse monoclonal anti-human EPCAM (Santa Cruz Biotechnology, Santa Cruz, CA, USA) antibody, followed bywith an anti-mouse Alexa 647 conjugated antibody (Abcam, Cambridge, UK), was used for epithelial cell detection. Cell fluorescence was analyzed with an SP5 Leica Confocal Microscope.

#### Inflammasome activation assessment

THP1-ASC-GFP were maintained in RPMI medium containing 10% FBS, 100U/mL penicillin and 100µg/mL streptomycin at 37°C in a 5% CO_2_ atmosphere and plated at 3.6×10^5^ cells/well in 96-well plates. Cells were stimulated with 0.5mM PAH/TPP NP or primed with ultrapure 1µg/mL LPS from *E. coli* followed with 0.5mM PAH/TPP NP ON. For a positive control, cells were primed with 1µg/mL LPS from *E. coli* for 3h and transfected with 500ng/mL bacterial plasmid cDNA. pcDNA was generated by cloning the PCR-generated full-length cDNA from a random non-related gene. Transfection was carried out with Lipofectamine^TM^ LTX Reagent with PLUS TM Reagent (Invitrogen, Thermo Fisher Scientific, Waltham, MA, USA), following the manufacturer’s instructions. Supernatants were harvested and stored at −80°C until used and cells were stained with DAPI (nuclear staining) (Invitrogen, Thermo Fisher Scientific, Waltham, MA, USA) and fixed with 2% PFA fixation for the microscopy analysis.

The frequency of ASC-GFP^+^ cells and localization of fluorescent ASC specks were assessed using a Nikon Eclipse Ti Fluorescence Microscope and analyzed with ImageJ software. The total cells: ASC-GFP^+^ cells ratio was calculated.

#### Quantification of secreted cytokines

Supernatants of stimulated BMDM, J774, BMDC, THP1, THP1-ASC-GFP and HT-29 cells were assessed for mIL-1β, mIL-18, mIL-6, mIFN-γ and mTNF-α using the Mouse DuoSet R&D Systems ELISA kit. The concentration of hIL-1β and hIL-8 were measured using a human Invitrogen ELISA Kit (Thermo Fisher Scientific, Waltham, MA, USA). All the protocols were used according to the manufacturer’s instructions. The concentration of released LDH (enzymatic assay) (LDH-L Weiner Lab, Rosario, Santa Fe, Argentina) was quantified.

#### Immunoblotting analysis

Cell lysates and supernatants were analyzed by SDS-PAGE followed by immunoblotting. Proteins suspended in loading buffer (SDS+ϕ3-mercaptoethanol) were separated on a 15% SDS-PAGE gel, and transferred onto nitrocellulose membranes (Amersham Biosciences, Uppsala, Sweden) using transfer buffer (50mM Tris, 40mM glycine, 10% methanol). After blocking for 1h in TBS with 0.1% Tween-20 containing 5% non-fat dry milk, membranes were incubated overnight at 4°C with a mouse monoclonal antibody to the p20 subunit of caspase-1 (Adipogen, San Diego, CA, USA) at a 1:1,000 dilution. A loading control blot was performed with an anti–β-actin monoclonal antibody (Cell Signaling Technology, Danvers, MA, USA) at a 1:1,000 dilution. Following three washes in TBS with 0.1% Tween 20, membranes were incubated for 1h at RT with the appropriate HRP-conjugated secondary antibody (Cell Signaling Technology, Danvers, MA, USA) at a 1:1,000 dilution. Immunoreactive bands were visualized using Luminol chemiluminescent HRP substrate (Millipore, Burlington, MA, USA) and analyzed with ImageQuant TL Software (GE Healthcare, Buckinghamshire, UK).

### Analysis of the LPS content of PAH/TPP nanoparticles

The HEK-hTLR4-reporter cell line was maintained in DMEM containing 10% FBS, 100U/mL penicillin and 100µg/mL streptomycin at 37°C with 5% CO_2_. Thirty thousand cells/well were seeded in 96-well microplates and incubated to 70% confluence. The medium was fully replaced and cells were stimulated ON with different concentrations of LPS, 0.5mM PAH, 0.3mM TPP, or 0.5mM PAH/TPP NP. The supernatants were harvested and the secreted embryonic alkaline phosphatase (SEAP) was measured.

### *In vivo* assessment of adjuvant properties

#### Inflammasome characterization

Male six- to nine-week-old wild-type and KO C57BL/6 Nlrp3^-/-^, Casp1/11^-/-^, Casp11^-/-^, Gsdmd^-/-^ and IL-1R^-/-^ mice were purchased from the Federal University of Minas Gerais and maintained under 12h cycles of light/dark. Femur and tibiae were removed for BMDM preparation and for immunization assays. Furthermore, male five- to seven-week-old wild-type Balb/c mice were purchased from the Laboratory of Experimental Animals, School of Veterinary Sciences of the University of La Plata (UNLP), Argentina.

Mice (C57BL/6 and Balb/c wild-type mice, and null C57BL/6 mice) were immunized once a week for three consecutive weeks intraperitoneally (IP) with OVA-conjugated PAH/TPP NP (NP-OVA) or OVA adjuvanted with aluminum hydroxide (OVA+Alum). As a control, mice were immunized only with 100µg OVA EndoFit^(TM)^. One week after the last immunization, mice were bled and sacrificed for the immune response analysis.

#### Humoral response assessment

OVA-specific antibodies were measured by indirect ELISA in different biological samples. Briefly, MaxiSorp plates (NUNC, Roskilde, Denmark) were coated with 1µg/mL OVA, 5mM PAH or 5mM NP in carbonate-bicarbonate buffer, pH=9.6. Coated plates were blocked with 5% equine serum in saline for 1h at 37°C, and subsequently incubated with sera (1/100) for 1h at 37°C. Bound immunoglobulins were detected with anti-mouse IgG-HRP (Biorad, Hercules, CA, USA), anti-mouse IgG1-HRP, anti-mouse IgG2a-HRP (BD Biosciences, Franklin Lakes, NJ, USA) or anti-mouse IgA-HRP (Thermo Fisher Scientific, Waltham, MA, USA). The reaction was developed with TMB (Sigma-Aldrich, St. Louis, MO, USA) and stopped with 2M H_2_SO_4_. OD was measured at 450 nm with Varioskan Lux (Thermo Fisher Scientific).

#### Assessment of secreted cytokine from stimulated splenocytes

Spleen cell suspensions were prepared in RPMI-1640 supplemented with 10% FBS, 10U/mL penicillin and 10mg/mL streptomycin, seeded (4×10^6^ cells/mL) in culture plates and cultured for 72h at 37°C under controlled atmosphere. Cells were recalled with 10μg/mL OVA, and supernatants were harvested for mIFN-γ, mIL-5 and mIL-13 quantification by ELISA, using commercially available kits (R&D Systems, Minneapolis, MN, USA) and following the manufacturer’s instructions.

Cytokines were also assessed by intracytoplasmic flow cytometry in spleen cells. Splenocytes were incubated with brefeldin A (BD Biosciences, Franklin Lakes, NJ, USA) for the last 4h of culture and stained with anti-CD4 (PerCP-Cy5.5) anti-CD8 (APC) monoclonal Ab (Thermo Fisher Scientific, Waltham, MA, USA). Cells were fixed, permeabilized, and stained with an anti-mouse IFN-γ (FITC) (Thermo Fisher Scientific, Waltham, MA, USA). Sample acquisition was performed with a FACS Aria Fusion cytometer, and data were analyzed using the FlowJo software.

#### Ethics

All experimental protocols were conducted in strict agreement with international ethical standards for animal experimentation (Helsinki Declaration and its amendments, Amsterdam Protocol of Welfare and Animal Protection and National Institutes of Health, USA NIH, guidelines: Guide for the Care and Use of Laboratory Animals). Anesthetized mice (isoflurane 5%) were killed by cervical dislocation by experienced research personnel, who performed it humanely and effectively. All efforts were made to alleviate suffering during the whole experiment. All protocols were approved by the Institutional Committee for the Care and Use of Laboratory Animals of the School of Sciences (University of La Plata) (Protocol Number: 006-37-21).

### Statistical analysis

All experiments were repeated at least twice with consistent outcomes. Flow cytometry data were analyzed using FlowJo X, images obtained by microscopy were analyzed using ImageJ. Graphs and data analysis were performed using GraphPad Prism8. Statistical analyses were carried out using one- or two-way ANOVA, followed by the Tukey test to discern significant differences among measurements from different experimental groups or by the t-Student test when applicable. Quantitative data are expressed as mean ± SEM, and statistical significance was defined as p<0.05.

## Supporting information

Rizzo et al. Supporting Information

## AUTHOR CONTRIBUTIONS

*GPR* Conceptualization, Data curation, Formal analysis, Investigation, Methodology, Visualization, Writing – original draft, Writing – review & editing. *RCS* Data curation, Methodology. *CC* Data curation, Formal analysis, Investigation, Methodology. *DSB* Data curation, Formal analysis, Investigation, Methodology. *EA* Data curation, Methodology. *SEH* Data curation, Methodology, Writing – original draft, Writing – review & editing. *MLA* Data curation, Methodology. MLC Data curation, Methodology, Writing – review & editing. WAM Data curation, Methodology, Writing – review & editing. IK Formal analysis, Methodology, Writing-review & editing. AT Formal analysis, Methodology, Writing-review & editing. *SCO* Supervision, Visualization, Writing – original draft, Writing – review & editing. *OA* Supervision, Visualization, Writing – original draft, Writing – review & editing. *PLS* Conceptualization, Data curation, Formal analysis, Funding acquisition, Investigation, Methodology, Supervision, Visualization, Writing – original draft, Writing – review & editing. *GHD* Conceptualization, Formal analysis, Funding acquisition, Investigation, Supervision, Visualization, Writing – original draft, Writing – review & editing.

## ACKNOWLEDGMENTS

SHE, MLA, MLC, WAM, AT, OA, PLS and GHD are researchers of CONICET; RCS and SCO are researchers of *Federal University of Minas Gerais; GPR, DSB and EA are* fellows of CONICET; CC and IK are fellows of ANPCyT.

