## Supplementary material for "Poly(allylamine)/tripolyphosphate nanocomplex coacervate as a NLRP3-dependent systemic adjuvant for vaccine development": Rizzo et al. Supporting Information

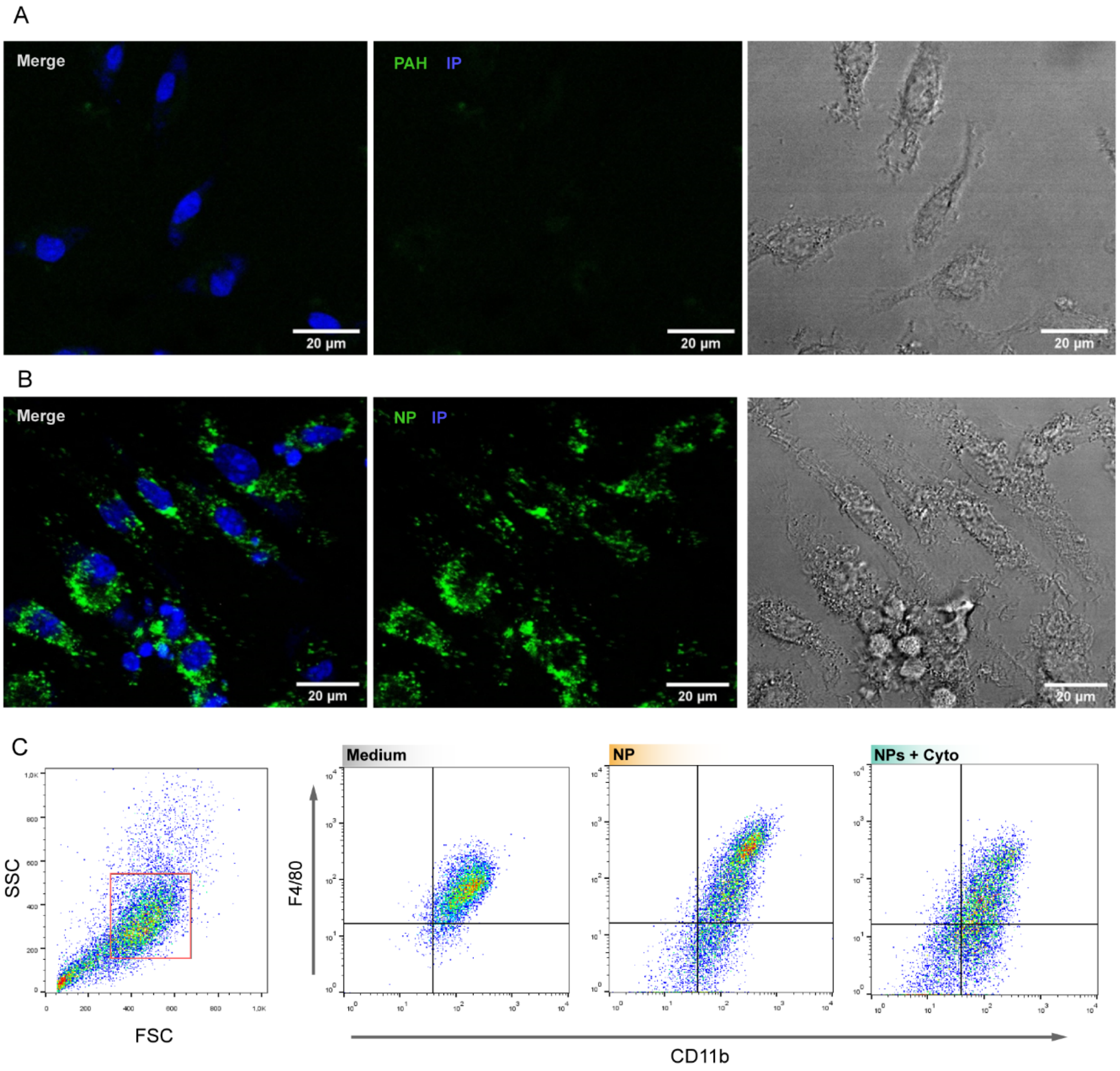


**Figure S1: Internalization of (A) PAH-FITC (B) and NP-FITC** in J774 macrophages by confocal microscopy. Representative images are shown. **(C)** Gate strategy to select F4/80 CD11b-positive macrophages cells.

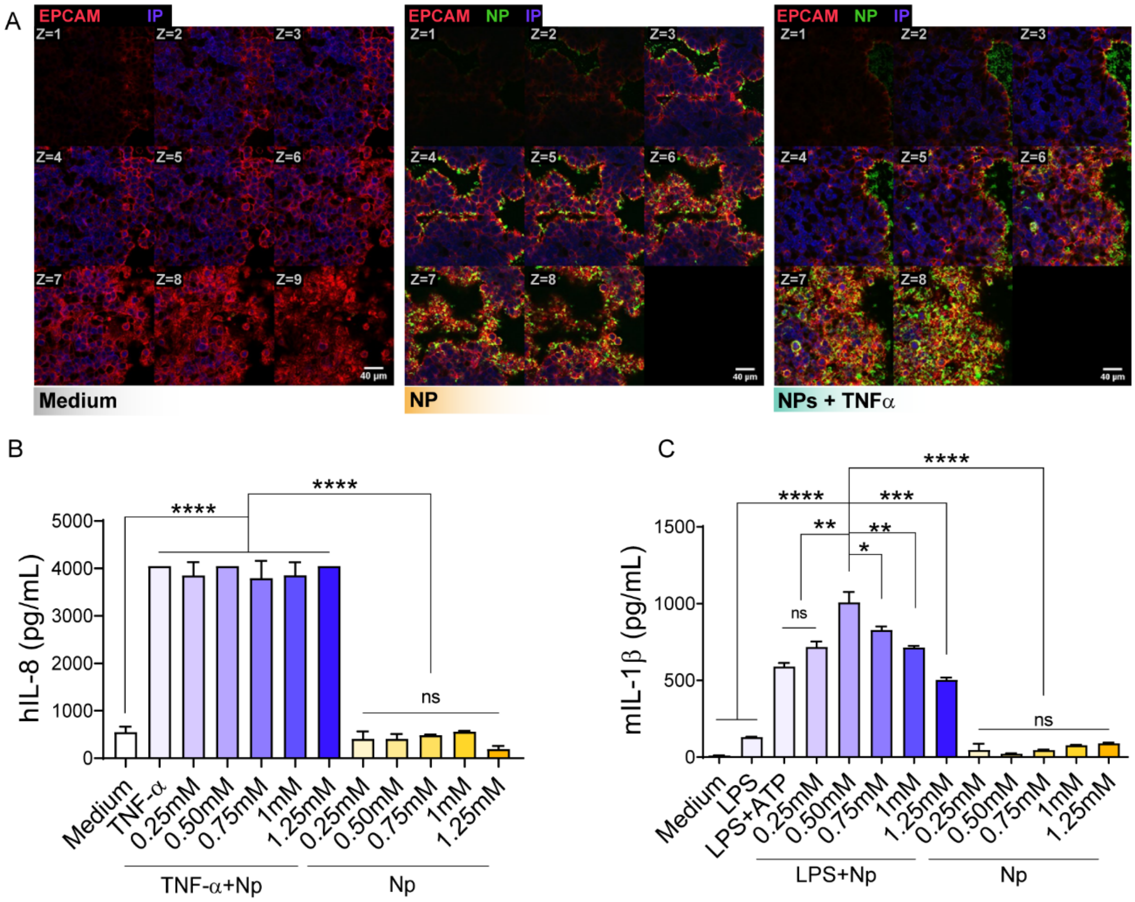


**Figure S2**: **Interaction of NP-FITC with human epithelial cell lines and dose-dependent cytokine response of cells to different stimuli.** **(A)** HT-29, different z-axis positions were scanned by confocal microscopy and EPCAM were used as a marker of epithelial cells. Representative images are shown. **(B)** Quantification of secreted hIL-8 in HT-29 cells exposed to medium, TNF-α, TNF-α+NP and NP; **(C)** Quantification of secreted mIL-1β in J774 macrophages exposed to medium, LPS, LPS+ATP, LPS+NP and NP. All experiments were carried out in triplicate and date are expressed as mean±SEM. P-value was determined by ordinary one-way ANOVA *p<0,05; **p<0,01; ***p<0,001; ****p<0,0001.

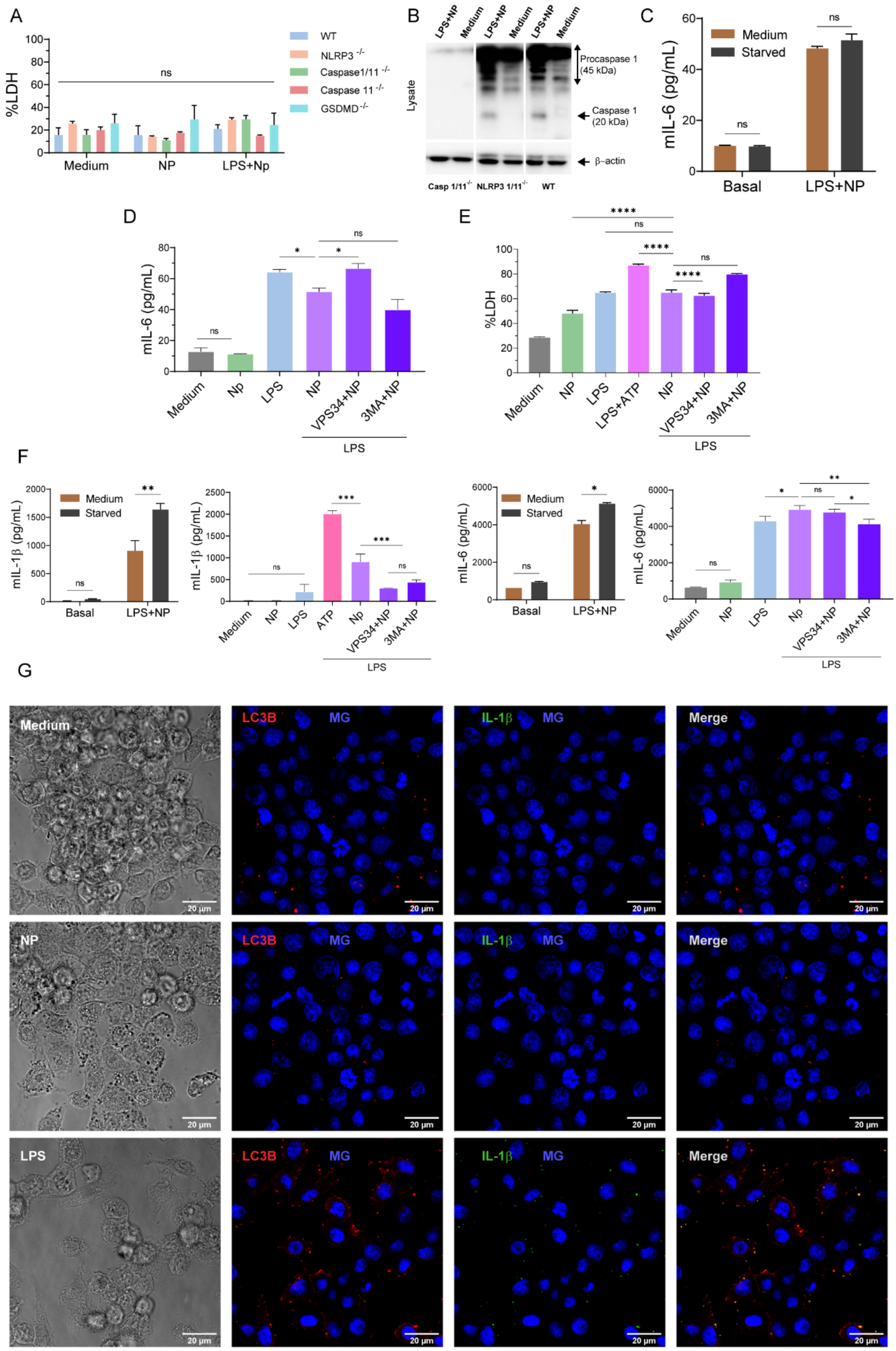

**Figure S3**: **Inflammasome pathway analysis in BMDM from C57BL/6 (WT), NLRP3$^{-/-}$, Casp1/11$^{-/-}$, Casp11$^{-/-}$, GSDMD$^{-/-}$ mice.** Cells were stimulated with medium, NP or LPS+NP, and **(A)** %LDH were quantified in the supernatants and **(B)** caspase-1 activation was analyzed by immunoblotting in cell lysates using an antibody specific for the p20 subunit (20kDa) or the procaspase-1 (45kDa). J774 cell lines remained unstimulated or were stimulated with LPS. Three hours later, they were treated with NP. At 6 hours post-NP stimulation, cells were either left untreated (in medium) or their supernatants were removed and replaced with EBSS for an additional hour of culture. Subsequently J774 macrophages were stimulated for 6h with NP in the presence of inhibitors of the autophagy VPS34-IN1 and 3-methiladenine (3-MA) and quantification **(C-D)** mIL-6 and **(E)** %LDH in the supernatants. **(F)** Quantification of mIL-1β and mIL-6 in the supernatants of BMDCs after the treatment with autophagy inhibitors. **(G)** Medium, NP and LPS controls for the mIL-1β-LC3B colocalization by confocal microscopy. P-value was determined by ordinary one-way ANOVA *p<0,05; **p<0,01; ***p<0,001; ****p<0,0001.

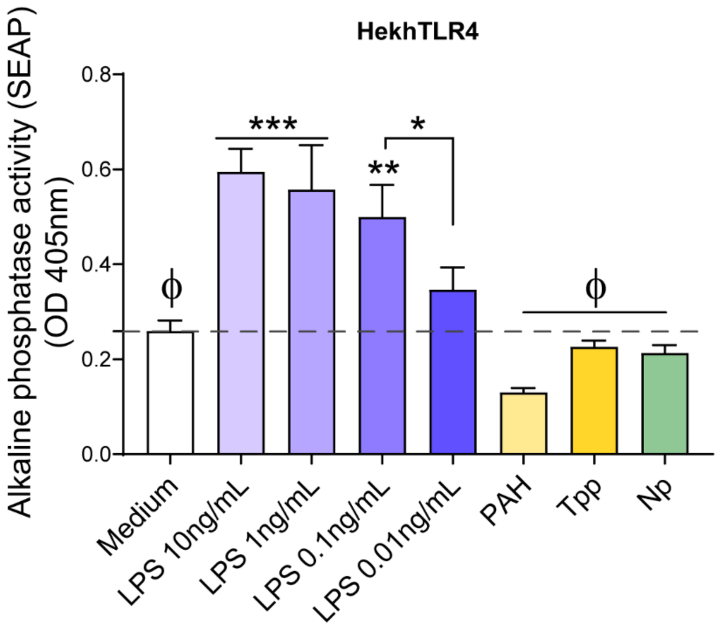


**Figure S4**: **Analysis of the stimulation of the Hek-hTLR4 cell line with LPS**. Different concentrations of LPS were included as positive controls; SEAP activity was measured in the supernatants. This experiment was performed in triplicate and data are expressed as mean±SEM. P-value was determined by ordinary one-way ANOVA *p<0,05; **p<0,01; ***p<0,001; Φ a result significantly different from LPS.

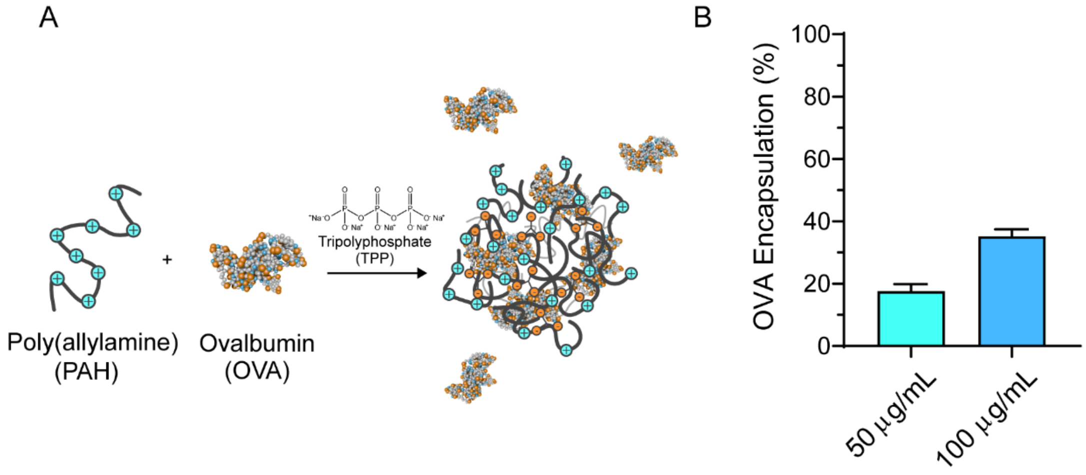


**Figure S5: Analysis of the OVA encapsulation by NPs. (A)** Scheme of protein uptake during PAH/TPP NP formation. As OVA present a negative net charge at the working pH, free proteins act as PAH ionic crosslinkers along with TPP anions. **(B)** OVA Loading percentage (%) analysis. Free OVA was analyzed in the supernatant of the PAH/OVA/TPP NP formulation by protein quantification (BCA). This experiment was performed in triplicate and data are expressed as mean±SEM. P-value was determined by Student's test ### p<0,001.
